# Sustained Vaccine Exposure Elicits More Rapid, Consistent, and Broad Humoral Immune Responses to Multivalent Influenza Vaccines

**DOI:** 10.1101/2024.04.28.591370

**Authors:** Olivia M. Saouaf, Ben S. Ou, Ye Eun Song, Joshua J. Carter, Jerry Yan, Carolyn K. Jons, Christopher O. Barnes, Eric A. Appel

## Abstract

With the ever-present threat of pandemics, it is imperative we develop vaccine technologies eliciting broad and durable immunity to high-risk pathogens. Yet, current annual influenza vaccines, for example, fail to provide robust immunity against the 3-4 homologous strains they contain, let alone heterologous strains. Herein, we demonstrate that sustained delivery of multivalent influenza vaccines from an injectable polymer-nanoparticle (PNP) hydrogel technology induces more rapid, consistent, and potent humoral immune responses against multiple homologous viruses, as well as potent responses against heterologous viruses and potential pandemic subtypes H5N1, H7N9 and H9N2. Further, admixing PNP hydrogels with commercial influenza vaccines results in stronger hemagglutination inhibition against both heterologous and homologous viruses. We show this enhanced potency and breadth arises from higher affinity antibodies targeting both the hemagglutinin stem and head. Overall, this simple and effective sustained delivery platform for multivalent annual influenza vaccines generates durable, potent, and remarkably broad immunity to influenza.

## 1. Introduction

Vaccine technologies are our strongest tools in combatting viral infections. Yet, despite their dramatic advancements over the past few years in the wake of the COVID-19 pandemic, current vaccine technologies are unable to eliminate threats posed by highly mutagenic viruses like influenza, which still causes roughly 500,000 respiratory deaths worldwide annually^[1]^. Over the past decade, seasonal influenza vaccines have had an average effectiveness of just 40%^[2]^, and influenza infection rates have quickly returned to pre-COVID-19 pandemic levels^[3]^. While typical seasonal influenza vaccines comprise the 3-4 subtypes predicted to be the most prevalent each year, the challenges with selecting the correct viruses to include each year are compounded by the dramatic variability in effectiveness against each influenza virus in the vaccine^[4–7]^. Additionally, the threat of interspecies transmission of highly pathogenic strains such as H5N1 avian influenza remains at the forefront of our disease monitoring as growing population density and large-scale agriculture increase the opportunities for viral mutation and zoonotic spillover^[8–10]^. Indeed, current influenza vaccines focus on predominantly human-to-human transmitted strains (e.g., H1N1, H3N2, B Victoria, and B Yamagata), leaving our population with little to no protection against a potential H5N1 pandemic outbreak. To combat currently human-circulating influenza viruses as well as preempt future interspecies pandemic strains, it is essential we improve our current influenza vaccine technologies to elicit more potent and durable responses against homologous viruses and enable efficacious responses against never-before encountered heterologous strains. The ideal influenza vaccine technology would therefore induce consistent seroconversion and high antibody titers against all strains covered by the vaccine, as well as robust breadth against future strains. Working towards these goals will bring us closer to a globally accessible universal influenza vaccine.

When innovating on the annual influenza vaccine, it is important to consider ease of scale up and distribution to a global market while still achieving the goal of generating more potent, broad, and durable immunity. Most influenza vaccines have historically comprised either inactivated or split viruses manufactured through propagation in chicken eggs, which can be slow and burdensome. Newer subunit vaccines are scalable, safe, and more easily stored than other vaccines as they use only the most relevant surface proteins of viruses of interest as a protein antigen to elicit immunity^[11]^. Each of these vaccine types supports the inclusion of immune activating adjuvants, which heighten the body’s immune response. For example, squalene oil-in-water emulsions such as MF-59 (Novartis) and AS03 (GlaxoSmithKline) have been shown to greatly improve vaccine effectiveness over non-adjuvanted formulations, but also exhibit more severe side effects^[12–14]^. Toll-like receptor (TLR) agonist adjuvants have shown exceptional promise as they enhance the activation and maturation of antigen presenting cells^[15–16]^. For example, CpG oligodeoxynucleotide is a TLR9 agonist that has been used in commercial vaccines (e.g., Hepislav-B; Dynavax) and Monophosphoryl lipid A (MPLA) is a TLR4 agonist that has been incorporated into several commercial adjuvants (e.g., AS04, part of Fendrix and Cervarix; GlaxoSmithKline), and both have been shown to be effective as adjuvants in influenza vaccines^[17–19]^. More recently, the potent TLR7/8 agonist 3M-052 (an imidazoquinoline derivative; 3M Corporation) has demonstrated exceptional promise in enhancing humoral immunity to HIV envelope proteins as well as SARS-CoV-2^[20–22]^. Yet, despite their promise, adjuvants alone have not successfully demonstrated the ability to generate sufficiently potent and broad immunity^[23–24]^.

Recent studies have demonstrated that precise spatiotemporal control of vaccine exposure greatly affects the magnitude and quality of resulting immune responses^[25–29]^. In the context of HIV vaccines, researchers have shown that extended vaccine delivery over the course of 2-4 weeks, either with surgically implanted osmostic pumps or by splitting a given vaccine dose into multiple injections^[26, 30–32]^, can increase the potency and breadth of humoral immune responses. Similarly, prolonging influenza vaccine exposure to the immune system by splitting a dose over weeks of daily injections yields higher quality immune responses^[31]^. Various studies have demonstrated that extended vaccine exposure prolongs stimulation of germinal centers (GCs) in the draining lymph nodes, enabling higher affinity maturation^[28–29, 33]^ and modulating immunodominance of certain epitopes^[30, 32]^, potentially generating more potent responses against a greater diversity of epitopes on a given antigen or against poorly immunogenic antigens^[34]^.

While influenza has two primary surface antigens, hemagglutinin (HA) and neuraminidase (NA), HA has been the focus of most antigen engineering efforts as it tends to dominate immune responses when delivered in conjunction with NA^[35–39]^, as in traditional inactivated or split virus vaccines. HA proteins have both a highly mutagenic and immunodominant head region (the outermost part of the influenza surface protein), and a stem region that remains highly conserved between various subtypes but exhibits poor immunogenicity^[40]^. Moreover, NA proteins have more highly conserved regions than HA proteins generally^[41]^. Extended delivery vaccination has the potential to generate more potent responses to the conserved HA stem region and/or NA protein, depending on the antigens used, thereby enhancing protection to a broader array of influenza strains. Unfortunately, burdensome repeat-dosing regimens and surgically implanted pumps are not feasible clinically, necessitating the development of technologies compatible with existing vaccine formats (e.g., inactivated virion or subunit vaccines).

To address this challenge, our lab has developed a self-assembled polymer-nanoparticle (PNP) hydrogel depot technology that is easily injectable^[42–43]^, highly biocompatible and non-immunogenic^[44]^, and which has been shown to both stabilize and sustain the delivery of numerous protein, cellular, and molecular therapeutic cargos^[45–51]^. We have previously shown this PNP hydrogel platform to be effective in providing robust and durable immune responses to numerous single-antigen vaccines for influenza and SARS-CoV-2^[52–58]^. In this work, we sought to leverage this hydrogel depot technology for the delivery of adjuvanted multivalent influenza vaccines to enhance the consistency, potency, and breadth of humoral immune responses. To demonstrate the ability of this technology to induce consistent immune responses across multiple antigens, we formulated a trivalent HA subunit influenza vaccine alongside MPLA in PNP hydrogels. We found that the PNP hydrogels achieved complete and robust seroconversion against all homologous strains and enhanced responses against heterologous strains compared to clinical adjuvant controls. Further, we then demonstrate the ability to simply admix a commercial quadrivalent influenza vaccine into a 3M-052-adjuvanted hydrogel, which enhanced the breadth of antibody titers and hemagglutination inhibition. We demonstrate that the observed enhancements in potency and breadth arise from higher affinity responses that target both the stem and head region of the HA antigen elicited by sustained vaccine exposure with the PNP hydrogels. In summary, this work demonstrates a simple and effective sustained delivery platform for multivalent annual influenza vaccines to create durable, potent, and remarkably broad immunity to influenza.

## 2. Results

### 2.1 Hydrogel for Slow Delivery of Influenza Vaccine

We have previously demonstrated that PNP hydrogels can reliably deliver single-antigen vaccines, prolonging GC reactions and resulting in improved humoral immune responses. Rapid formation of these hydrogels is achieved through the mixing of aqueous solutions of dodecyl-modified hydroxypropylmethylcellulose (HPMC-C_12_) with biodegradable nanoparticles (NPs) made from poly(ethylene glycol)-block-poly(lactic acid) (Figure 1a). The NPs act as physical crosslinkers of the network as HPMC-C_12_ polymers adsorb to the particles by strong entropy-driven, multivalent, and dynamic interactions^[59–61]^. In this work, we sought to leverage this extended-release drug delivery platform to generate potent and broad immune responses to multivalent influenza vaccines consisting of distinct strains of HA and a molecular adjuvant. Based on previous research^[52]^, we selected one PNP hydrogel formulation (2% HPMC-C_12_, 10% NP, 88% PBS/soluble vaccine components by weight; denoted PNP-2-10) to evaluate initially as a delivery vehicle in comparison to relevant liquid bolus controls. Our self-assembling PNP hydrogels lend themselves to delivery of various drug cargos as they are readily formed by simple mixing, they shear-thin through a needle for easy injection, and then rapidly self-heal into a robust solid-like depot in the body for sustained cargo delivery (Supplemental Figure 1). These properties allow for either fully formulated vaccine-loaded hydrogels to be preloaded into syringes, or for plain hydrogels to be admixed with liquid vaccines just prior to administration. In both cases, hydrogels must retain structural integrity under conditions for storage (5°C), administration at room temperature (25°C), and *in vivo* efficacy (37°C) (Figure 1b), so rheological amplitude sweeps were performed at each of these three temperatures (Figure 1c). Near constant overlap of the generated curves demonstrated that the solid-like properties of the PNP hydrogels are consistent at all relevant temperatures, illustrated by matching storage moduli at a selected strain amplitude (100%; Figure 1d). Additionally, the crossover of storage and loss moduli at near identical strain amplitudes shows that the PNP hydrogel yields similarly under all conditions tested, indicating injectability across the range of temperatures a clinician might encounter after removing a PNP-hydrogel based vaccine from refrigeration. These behaviors are consistent with the nature of the entropy-driven polymer-nanoparticle interactions^[60]^. To further verify injectability, we conducted injection force measurements on both PNP-1-5 and PNP-2-10 formulations through 12 mm long, 27 gauge ultra-thin-wall needles (equivalent in inner diameter to a standard 25 gauge needle) (Figure 1e). Both formulations were smoothly injected with no breakaway force and a glide force well below the limit for comfortable injection, indicating that our PNP hydrogel platform is easily usable in the clinic.^[43, 62]^ After injection, vaccine cargos have limited diffusivity within the dynamic hydrogel, allowing for their retention within the hydrogel depot over an extended period of time^[53]^. We performed a static release assay to evaluate cargo retention, placing our PNP hydrogel carrying fluorescently tagged albumin (protein of similar size to influenza hemagglutinin) and 3M-052 (TLR7/8 agonist adjuvant molecule) in a capillary tube and measuring cargo release into PBS (Figure 1f). Through this study we observed comparable, almost complete retention of the protein and small molecule cargo, indicating the ability for controlled co-delivery of the vaccine components (Figure 1g). 3M-052 was chosen as the model small molecule for this release study because it is the smallest of the three adjuvants used in this work and would therefore be expected to exhibit the fastest release time, providing a lower limit for adjuvant release time. This capillary release setup minimizes erosion of the hydrogel through small surface area contact with surrounding liquid, whereas the hydrogel depot is known to gradual dissolve over time *in vivo*, ensuring consistent release of entrapped cargo over time^[33, 48]^.

**Figure 1.**
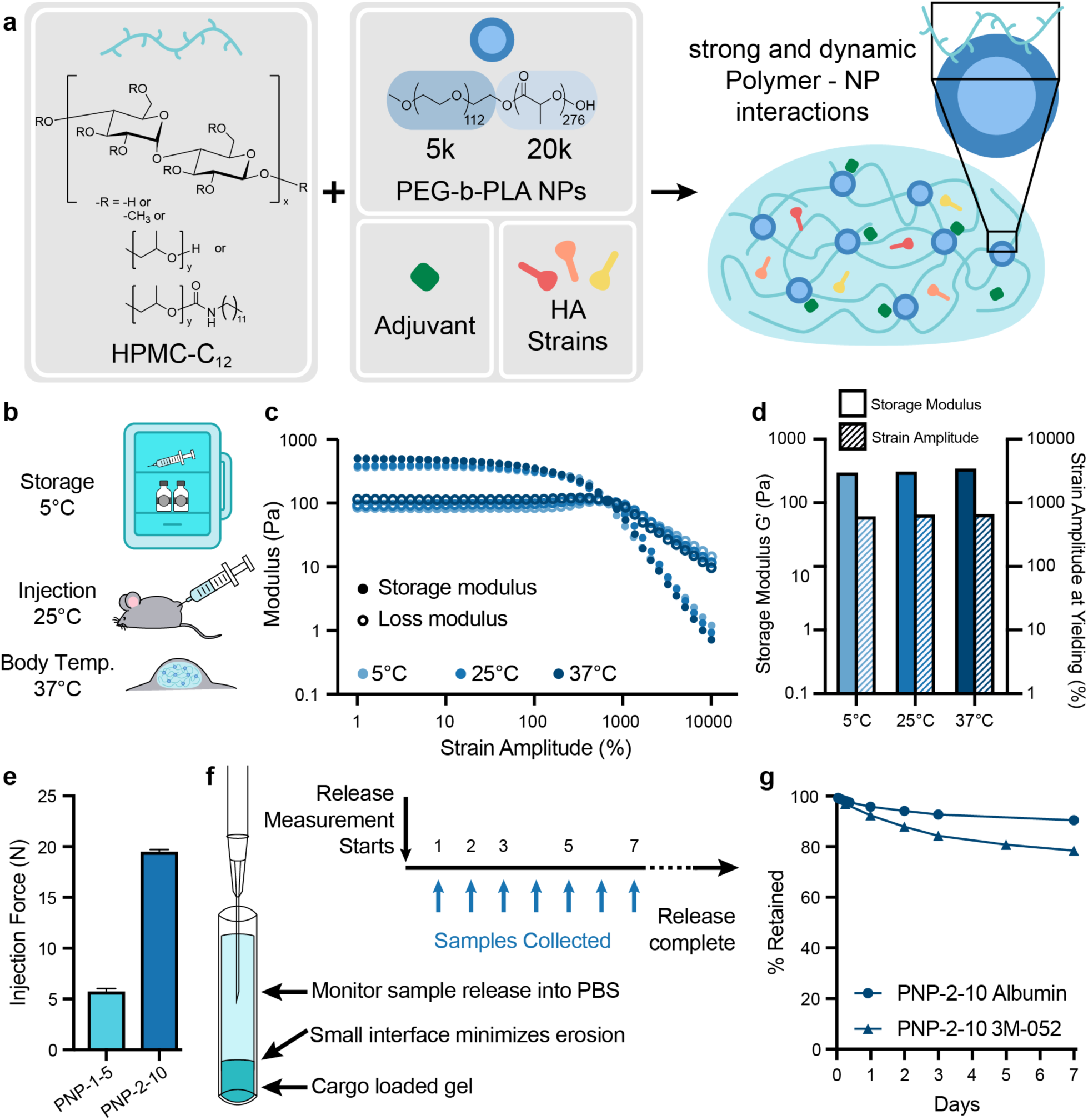
Multivalent influenza vaccines formulated in polymer-nanoparticle (PNP) hydrogels. **a.** Schematic of the mixing of HPMC-C_12_ with PEG-PLA NPs and vaccine cargoes (molecular adjuvants and antigens from multiple influenza strains) to form a vaccine-loaded PNP hydrogel. **b.** Representation of the temperatures at which a PNP-hydrogel based vaccine must function clinically. **c.** Stress-controlled oscillatory amplitude sweep rheometric assays performed on PNP-2-10 hydrogels at storage (5°C), room (25°C), and *in vivo* (37°C) temperatures. **d.** Plot of storage modulus at a strain amplitude of 100% and strain amplitude at yielding, defined as crossover between storage and loss moduli. **e.** Injection force of various PNP hydrogel formulations through a 12 mm long, 27 gauge ultra-thin-walled needle (n = 3). **f.** Diagram of a capillary release assay for evaluating release of cargo from PNP hydrogels into PBS and the associated sample collection schedule (n = 3). **g.** Retention kinetics of Alexa Fluor 647 BSA (M_W_ = 66 kDa, similarly sized to HA, M_W_ ≈ 60 kDa) (Albumin) and 3M-052 from PNP-2-10 in a glass capillary release assay. Data are shown as mean ± SEM.

In addition to the capillary release assay, we further examined the sustained release of protein cargo from the PNP hydrogels by conducting an *in vivo* HA release study (Figure 2). Mice (SKH1E, n = 5) were subcutaneously injected with with 5 µg fluorescently tagged HA protein (AF647-HA) from Influenza A H3N2 (A/Perth/16/2009) in 100 µL of either a PNP-2-10/MPLA hydrogel or an AS04-like Alum/MPLA adjuvanted liquid bolus. Persistence of the HA protein at the site of injection was evaluated using an *in vivo* imaging system (IVIS) using bright-field photographic imaging via standard camera overlaid with fluorescent imaging over 25 days (Figure 2a). AF647-HA was nearly undetectable at day 15 when administered in bolus but continued to persist in the PNP hydrogel depot beyond day 25 (Figure 2b). To evaluate total antigen persistence, we evaluated the area under the release curve and found that HA in PNP-2-10 had a significantly higher presence in the persistence timeframe when administered in PNP as compared to bolus administration (Figure 2c). The PNP-2-10 vehicle significantly increased t_80%_, time to 80% release (Figure 2d). These data show that the PNP platform extends the release of HA vaccine cargos as compared to a liquid bolus injection.

**Figure 2.**
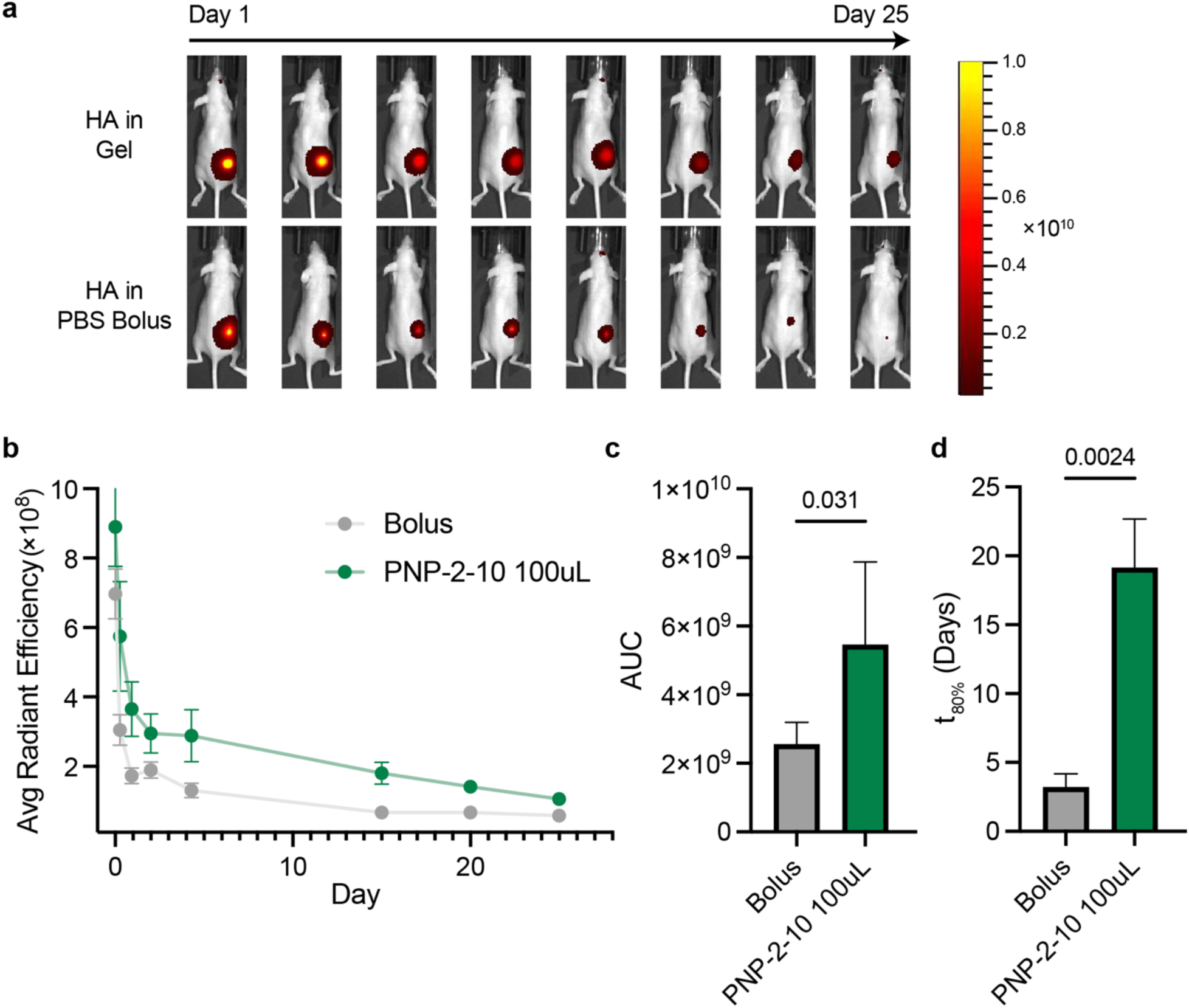
In vivo evaluation of hemagglutinin persistence upon subcutaneous vaccination. Mice were immunized (100 µL administration) with AF647-HA (5 µg) and MPLA (20 µg) formulated in PNP-2-10 or with alum in a liquid bolus. **a.** Representative images showing the different persistence times of HA from subcutaneous hydrogel or liquid bolus vaccination over 25 days. Units of fluorescence represented by the color bar are (p/sec/cm2/sr) / (µW/cm2). **b.** Radiant efficiency recorded over 25 days. **c.** Area under the curve of the radiant efficiency plot in (b). **d.** Time to 80% release (t_80%_) as calculated using a one-phase decay. Data are shown as mean ± SEM (n=5), and P values determined by a two-tailed Student’s t-test. Statistical significance was considered as P < 0.05.

### 2.2 Homologous Responses to a Trivalent HA Vaccine

*In vivo* experiments were then performed to evaluate the impact of slow vaccine delivery from our PNP hydrogel platform in eliciting humoral responses to a trivalent influenza subunit vaccine. Mice (C57BL/6) were immunized with a single dose of trivalent influenza vaccine containing 5 µg each of three hemagglutinin proteins (HA): Influenza A H1N1 (A/California/07/2009), Influenza A H3N2 (A/Perth/16/2009), and Influenza B (B/Brisbane/60/2008) (Figure 3a). Strains were selected to mimic clinical trivalent influenza vaccines which contain two influenza A subtypes – an H1N1 and an H3N2 – and one influenza B subtype (current quadrivalent clinical influenza vaccines contain both the Victoria and Yamagata lineage B subtypes, but this study used Victoria only). A single-dose regimen was utilized because the standard clinical influenza vaccine is administered as one shot annually. We prepared five different vaccine formulations: (i) Addavax (MF59-like squalene-based oil-in-water emulsion) adjuvanted liquid bolus, (ii) AS04-like Alum/MPLA adjuvanted liquid bolus, (iii) PNP-2-10/MPLA hydrogel, (iv) Alum/CpG adjuvanted liquid bolus, and (v) PNP-2-10/CpG hydrogel. Addavax was used as a clinically relevant representation of the current state-of-the-art for influenza vaccines. Liquid formulations including MPLA and CpG allowed evaluation of a novel adjuvant in the standard liquid form, and gel formulations containing MPLA and CpG were comparable to their respective alum-based counterparts to evaluate the effect of slow delivery from an immunological niche in conjunction with a novel molecular adjuvant. Sera were collected over a period of several months and IgG titers against homologous strains were evaluated via ELISA.

**Figure 3.**
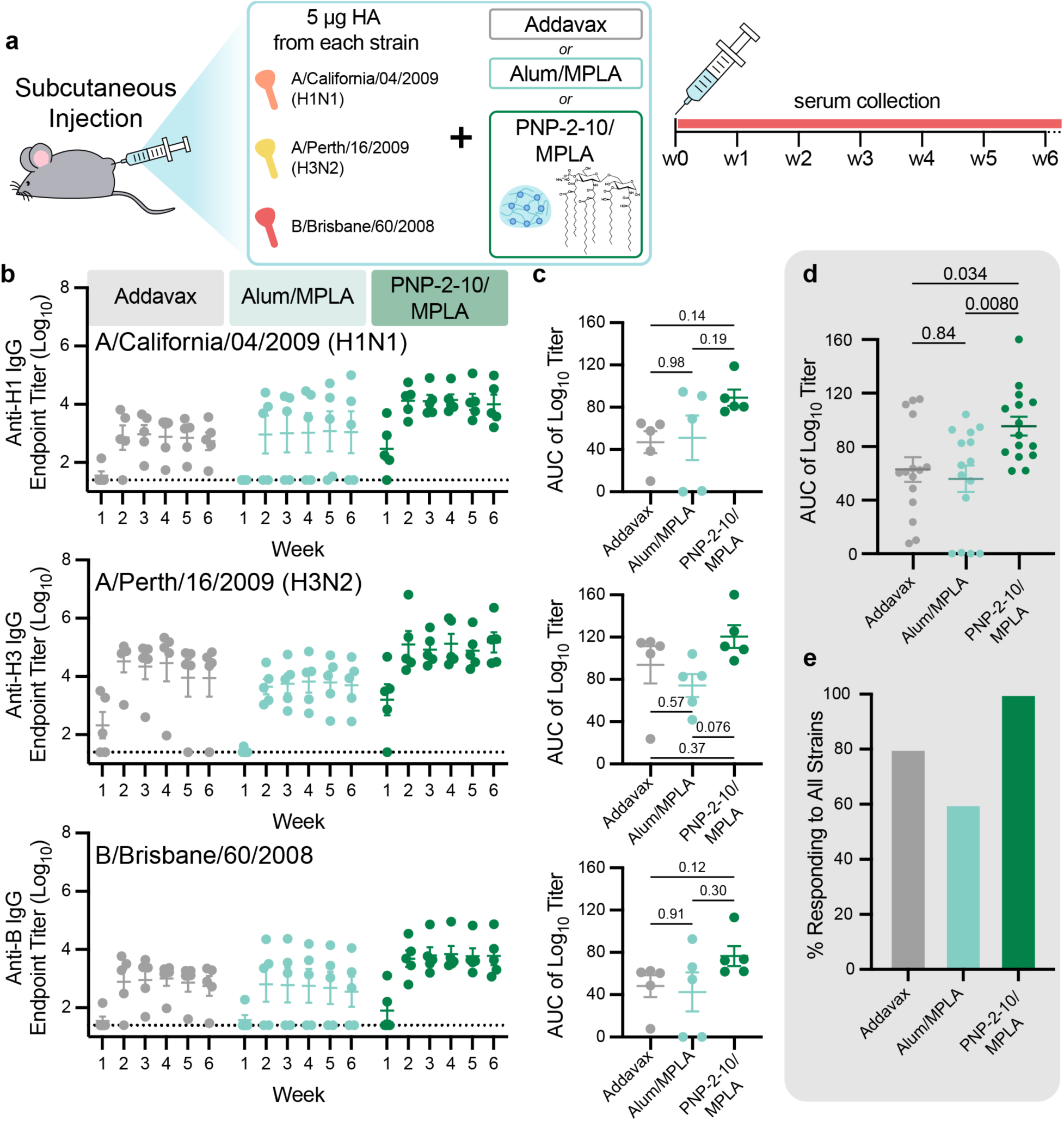
Trivalent subunit influenza vaccine adjuvanted by PNP hydrogels. Sustained delivery of multivalent flu vaccines induce robust, consistent humoral responses against all homologous strains. **a.** Trivalent subunit influenza vaccine schematic and *in vivo* vaccination/monitoring schedule. Mice were vaccinated with 5 μg HA protein from each strain (H1N1 (A/California/07/2009), H3N2 (A/Perth/16/2009), B (B/Brisbane/60/2008)) and an adjuvant (liquid Addavax, liquid Alum/MPLA, or PNP-2-10/MPLA) (n = 5). **b.** Anti-HA IgG titers from vaccinated mice over 6 weeks. Dotted lines indicate lower limit of detection. **c.** Area under the curve generated from the titers in (b). **d.** Area under the curve pooled across all HA strains. **e.** Percent of mice in each group responding to all strains, response defined AUC > 12. Data are shown as mean ± SEM, P values determined by one-way ANOVA with Tukey’s multiple comparisons test. Statistical significance was considered as P < 0.05.

Analysis of total IgG antibody titers against the three homologous strains used for vaccination showed that the PNP-2-10 group consistently outperforms bolus liquid vaccines (Figure 3, Supplemental Figure 3). Vaccines adjuvanted with MPLA were selected for further study because of more consistent performance as compared to CpG groups, but trends between alum and gel formulations were similar between the two molecular adjuvants. The observed differences in responses elicited by the two adjuvants are likely due to differences in the TLR biology of the distinct pathways activated by these two different molecules (TLR4 for MPLA and TLR9 for CpG). A time-course across the evaluated timepoints showed that all mice in the PNP-2-10/MPLA group seroconverted against all three HA’s while standard liquid bolus groups contained non-responders and/or mice whose antibody titers dwindled soon after vaccination (Figure 3b). An evaluation of the area-under-the-curve (AUC) of the titer time-course demonstrates that the overall IgG antibody coverage level for mice immunized with PNP hydrogel-based vaccines was consistently higher than bolus groups (Figure 3c). Analysis of titer AUC combined over all homologous strains reveals that the PNP-2-10/MPLA group significantly outperforms Addavax and Alum/MPLA bolus groups (Figure 3d). We observed that 100% of all mice in the PNP-2-10/MPLA group responded to all three strains of the trivalent vaccine, while only 80% and 60% of bolus groups responded (Figure 3e).

### 2.3 Breadth of Responses to a Trivalent HA Vaccine

We then evaluated the anti-HA IgG endpoint titers for week 6 sera against heterologous influenza viruses to which the murine model had never been exposed (Figure 4, Supplemental Figure 3). PNP hydrogel vaccines induced significantly increased antibody titers against viruses of the same subtype – H1 (Figure 4a), H3 (Figure 4b), B (Figure 4c). In these experiments, we see that most liquid bolus vaccinated mice elicited no response to the heterologous strains, leaving them completely unprotected against strains not predicted to be dominant by vaccine designers, which is the leading cause of ineffectiveness of the annual flu vaccine. In contrast, most of the PNP-2-10/MPLA vaccinated mice showed antibody responses to the heterologous strains, even to strains from almost a decade into the future, indicating broader immunity to influenza. We next evaluated immune responses against other influenza subtypes not represented in the vaccine, including H5, H7, and H9 strains that are of high priority for zoonotic spillover and pandemic risk (Figure 4d). Once again, PNP-2-10/MPLA vaccinated mice showed complete responses with titers against these heterologous subtypes comparable to those observed for the liquid bolus groups against the homologous strains. The breadth of humoral immunity observed in PNP-2-10/MPLA vaccinated mice against all strains indicated that these hydrogel-based vaccines significantly outperformed the liquid bolus controls (Figure 4e). A petal plot of all titers gathered against homologous and heterologous strains makes apparent the greatly increased efficacy of PNP vaccine in inducing higher antibody titers (Figure 4f). Further, the phylogenetic tree of the HA strains examined in this study demonstrates that the heterologous proteins (especially H5, H7, and H9), were highly genetically distinct from the ones used for vaccination (Figure 4g). These combined findings indicate that the extended vaccine exposure time enabled by the PNP hydrogel depot greatly increased the diversity of antibody response.

**Figure 4.**
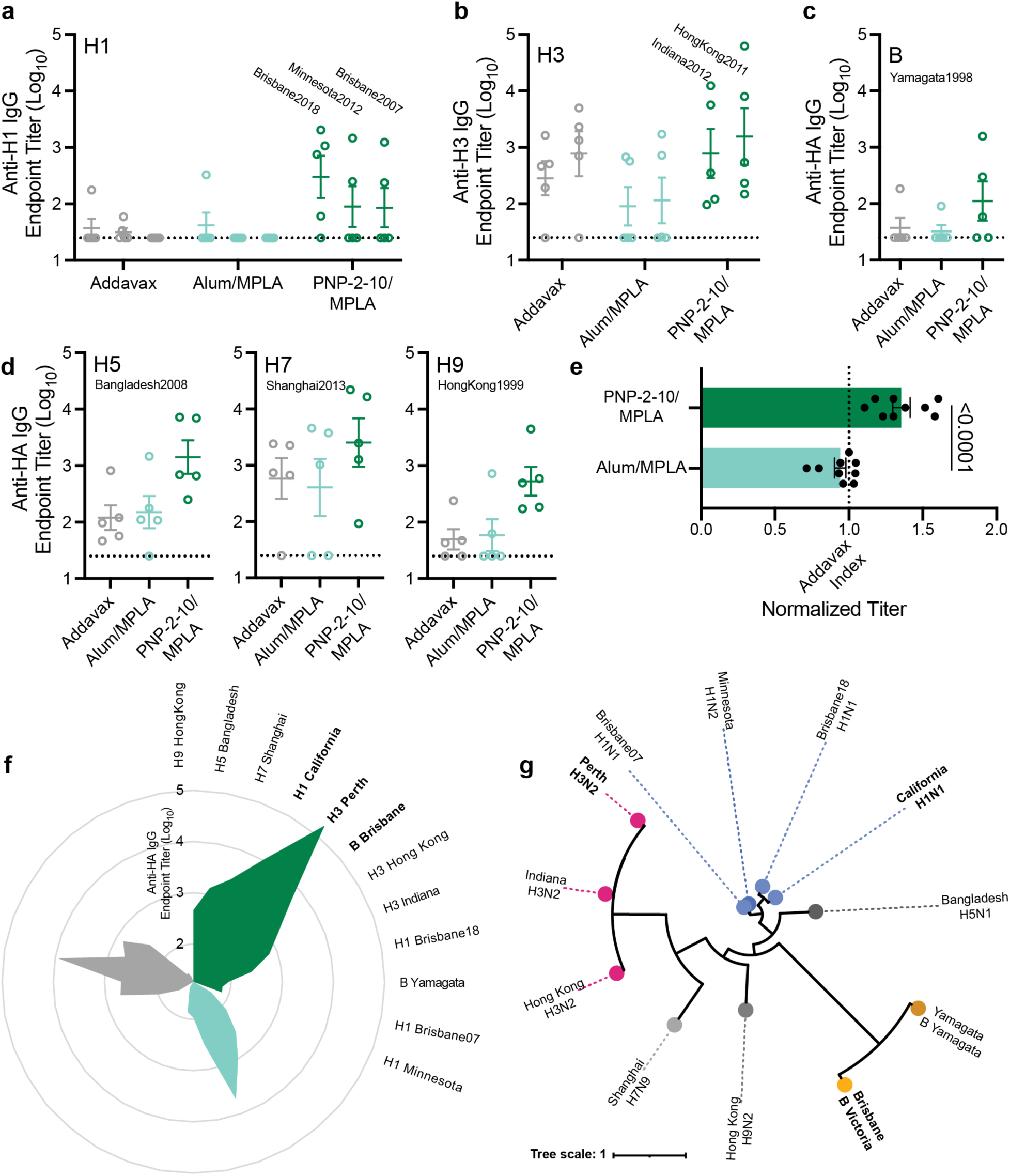
PNP hydrogel vaccine increases breadth of HA titers against heterologous strains. **a-c.** Week 6 IgG titers against HA from heterologous viruses bearing the same HA subtype – (a) H1, (b) H3 – or type – (c) B – as the HAs included in the administered vaccine. **d.** Week 6 Anti-HA IgG titers against H5, H7, and H9 -viral strains of different subtypes than the HAs included in the administered vaccine. Dotted lines indicate lower limit of detection. **e.** Average titer (log_10_) against all heterologous HA strains tested (a-d) normalized by the average Addavax titer (log_10_) against that same HA (dotted line Addavax Index). **f.** Petal plot summarizing average week 6 anti-HA IgG titers against all tested HAs (clockwise from top right: dark green PNP-2-10/MPLA, light green Alum/MPLA, gray Addavax). **g.** Phylogenetic tree of A/California/07/2009 (H1N1), A/Perth/16/2009 (H3N2), B/Brisbane/60/2008 (B Victoria), A/Brisbane/59/2007 (H1N1), A/Minnesota/14/2012 (H1N2), A/Brisbane/02/2018 (H1N1), A/Indiana/05/2012 (H3N2), A/Hong Kong/1514A01704826T/2011 (H3N2), B/Yamagata/16/1988 (B Yamagata), A/Bangladesh/207095/2008 (H5N1), A/Shanghai/MH01/2013 (H7N9), A/Hong Kong/1073/99 (H9N2) influenza antigens based on HA DNA sequence where the radial branch lengths represent modifications per site. Bold text indicates homologous strains. Created using the ‘Viral Genome Tree’ tool on the Bacterial and Viral Bioinformatics Resource Center (BV-BRC) website https://www.bv-brc.org/ and formatted on https://itol.embl.de/. n = 5 for all groups. All error bars are mean ± SEM, P value determined using a two-tailed Student’s *t*-test. Statistical significance was considered as P < 0.05.

### 2.4 Hydrogel Incorporation of a Clinical Quadrivalent Influenza Vaccine

Upon realizing the efficacy of our PNP hydrogel platform in enhancing the potency and breadth of a trivalent homemade subunit influenza vaccine, we chose to investigate its use in enhancing a widely clinically used inactivated influenza vaccine, Fluzone Quadrivalent (Figure 5). This off-the-shelf vaccine suited our purposes well: Fluzone is manufactured at two doses – high dose (0.086 mg/mL each HA strain), intended for the elderly population, and standard dose (0.03 mg/mL each HA strain) – each containing four strains of inactivated influenza (A/Victoria/2570/2019 (H1N1), A/Darwin/9/2021 (H3N2), B/Phuket/3073/2013, B/Austria/1359417/2021). The high dose can easily be admixed with our pre-fabricated adjuvant-containing PNP hydrogel, yielding a gel formulation standard dose (Figure 5a) with the same HA content and concentration as the clinical standard dose liquid (Figure 5b). The adjuvant 3M-052, a TLR7/8 agonist, was selected for inclusion in this formulation based on our previous work demonstrating that sustained co-delivery of TLR7/8 ligands by conjugation to the hydrogel network yielded enhanced humoral responses to vaccine and immunotherapy cargos^[53, 63]^. The lipidated 3M-052 adheres well to our NPs similarly to other lipidated molecules and can be simply formulated into PNP hydrogels^[48]^. Frequency sweep rheology shows that PNP-1-5 hydrogel formulated conventionally or via admixing maintained the same mechanical properties (Figure 5c, Supplemental Figure 4).

**Figure 5.**
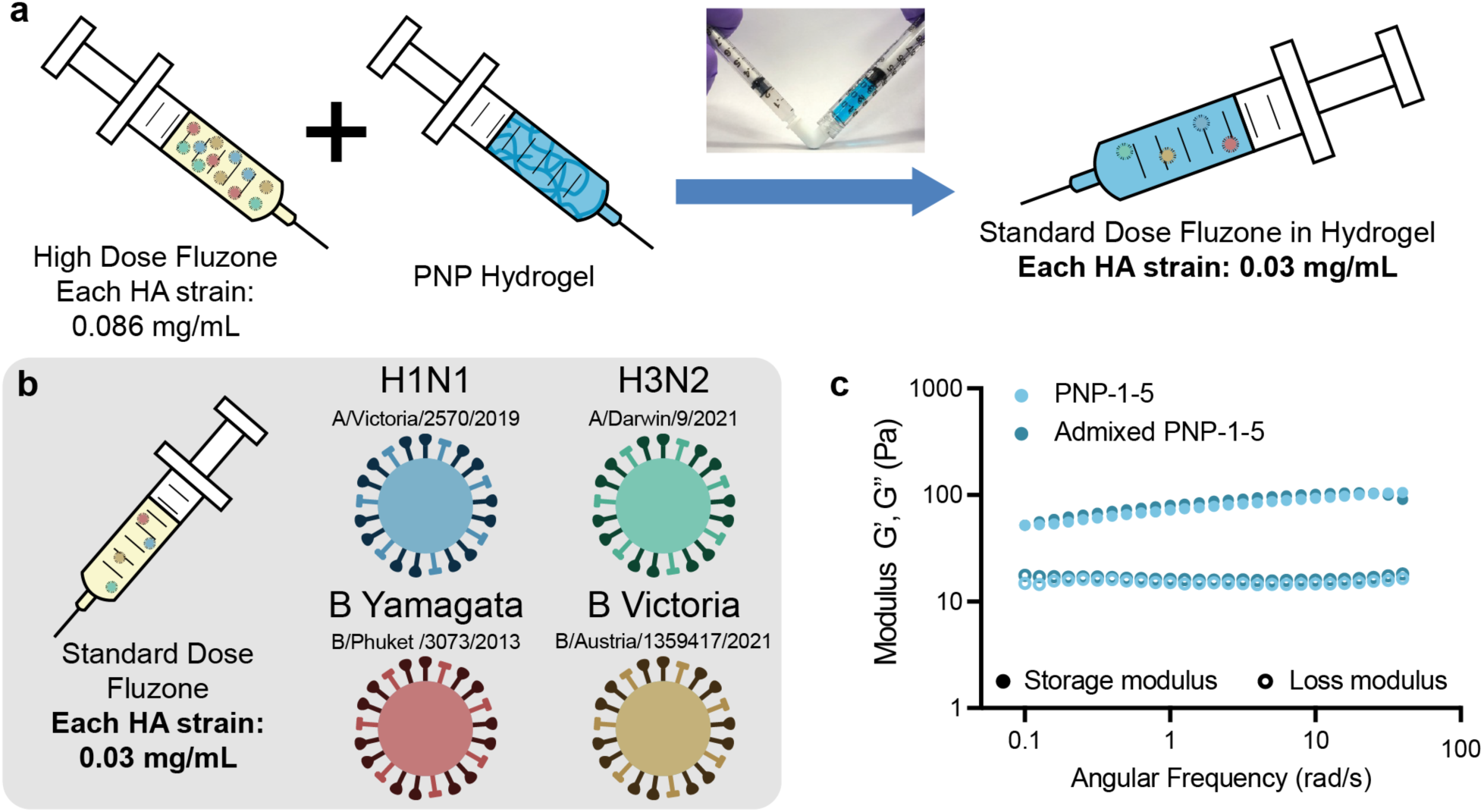
Incorporation of the clinical Fluzone Quadrivalent vaccine into PNP hydrogel. **a.** Schematic of mixing Fluzone High-Dose Quadrivalent with a concentrated PNP hydrogel via elbow mixer, resulting in a standard dose hydrogel formulation. **b.** Depiction of the standard dose Fluzone Quadrivalent vaccine containing four strains of inactivated influenza virus, each at a HA concentration of 0.03 mg/mL. **c.** Frequency sweep rheology of a conventionally made PNP-1-5 and an admixed PNP-1-5 (concentrated hydrogel made first and then diluted via elbow mixer).

### 2.5 Homologous Responses to a Clinical Quadrivalent Vaccine

To evaluate these clinical vaccine formulations, *in vivo* experiments were again performed. Mice (C57BL/6) were subcutaneously immunized with a single dose of commercial quadrivalent 2022-2023 season Fluzone influenza vaccine containing 2.5 µg each of the four listed hemagglutinin proteins (Figure 5a). We prepared four different vaccines groups: commercial Fluzone, PNP-1-5/Fluzone, PNP-1-5/3M-052/Fluzone, and PNP-1-5/MPLA/Fluzone. Injection volumes for this study were increased to 200 μL to better mimic typical vaccine administration volumes used clinically in human children (250 μL) while accounting for the subcutaneous administration limits in mice. As larger hydrogel depots take longer to dissolve and deliver their cargo, we elected to use a lower solids-content PNP-1-5 hydrogel formulation to provide a similar overall release timeframe as 100 μL of PNP-2-10 hydrogels.

For each of the four homologous influenza strains, adjuvanted gel groups significantly outperformed the commercial liquid bolus control (Figure 6, Supplemental Figure 5). As 3M-052 was designed for high potency adjuvanticity, its incorporation elicited further increased titers compared to MPLA, so we focused our comparison on Fluzone, PNP-1-5/Fluzone, and PNP-1-5/3M-052/Fluzone. Though PNP-1-5/Fluzone demonstrated improvement over bolus Fluzone, these results were not statistically significant, likely due to the complexity of creating a strong immune response to 4 different strains of influenza and the absence of the formation of an inflammatory niche within the hydrogel depot. The timecourse across the four months of sera collection showed consistent seroconversion against homologous HAs for all immunized mice in the PNP-1-5/3M-052/Fluzone group at higher titers than both commercial Fluzone and PNP-1-5/Fluzone groups across all timepoints (Figure 6b). The commercial vaccine group had at least one mouse whose IgG titer fell below the limit of detection over the course of the 4 months, while mice vaccinated with PNP-1-5/3M-052/Fluzone retained detectable anti-HA IgG concentrations over the entire study. PNP-1-5/3M-052/Fluzone elicited higher overall IgG coverage, exhibiting IgG AUC values significantly higher than commercial Fluzone across all strains (Figure 6c).

**Figure 6.**
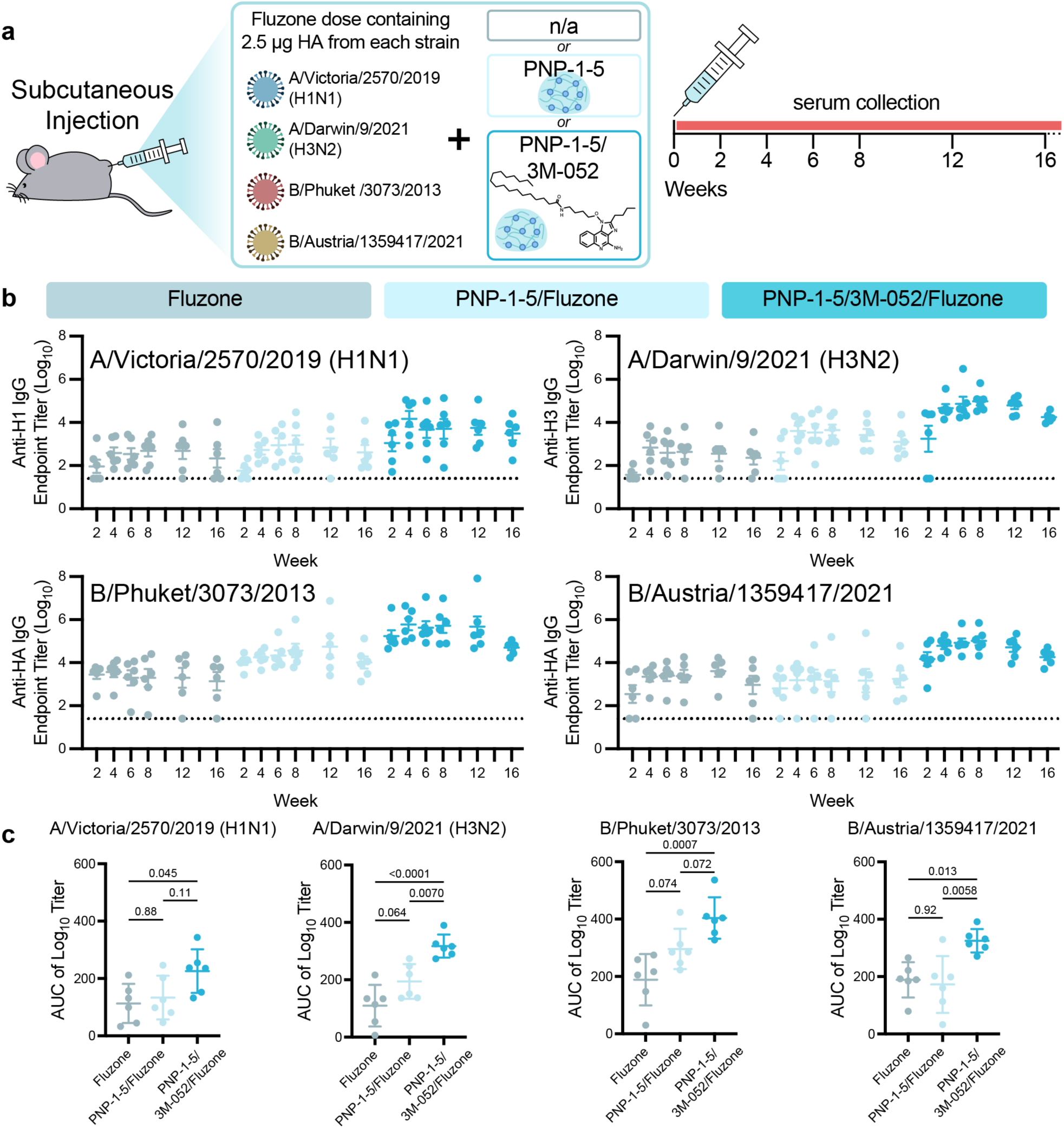
Admix of clinical quadrivalent Fluzone vaccine with adjuvanted PNP hydrogels induces robust, consistent humoral responses against all homologous strains. **a.** Quadrivalent inactivated influenza Fluzone vaccine schematic and *in vivo* vaccination/monitoring schedule. Mice were vaccinated with doses containing 2.5 μg HA protein from each strain of the 2022-2023 Fluzone Quadrivalent vaccine (H1N1 (A/Victoria/2570/2019), H3N2 (A/Darwin/9/2021), B/Yamagata lineage (B/Phuket/3073/2013), B/Victoria lineage (B/Austria/1359417/2021) and an optional adjuvant (PNP-1-5 or PNP-1-5/3M-052) (n = 6). **b.** Anti-HA IgG titers from vaccinated mice over 16 weeks. Dotted lines indicate lower limit of detection. **c.** Area under the curve generated from the titers shown in (b). All error bars are mean ± SEM, P values determined by one-way ANOVA with Tukey’s multiple comparisons test. Statistical significance was considered as P < 0.05.

### 2.6 Breadth of Immune Responses to a Clinical Quadrivalent Vaccine

In addition to quantifying homologous anti-HA titers, we examined humoral responses to heterologous HA and the less immunogenic but more highly conserved homologous NA antigens (Figure 7a). Anti-HA ELISAs were performed at week 6 to evaluate titers against heterologous influenza strains (Figure 7b). PNP-1-5/3M-052/Fluzone significantly increased titers from all other groups. Notably, nearly all members of Fluzone and PNP-1-5/Fluzone groups had no response to heterologous H1, whereas the PNP-1-5/3M-052/Fluzone group increased responders to >50%. In the cases of heterologous H3 and B strains, PNP-1-5/3M-052/Fluzone exhibited mean titers that were over an order of magnitude higher than those from commercial Fluzone. Homologous anti-NA ELISAs were performed at week 8 (Figure 7c). The PNP-1-5/3M-052/Fluzone group again demonstrated the highest titers, here against the more-conserved surface protein of influenza. Throughout this study, the PNP-1-5/Fluzone group had higher titers than commercial Fluzone, and the addition of the molecular adjuvant in PNP-1-5/3M-052/Fluzone further increased IgG response. A petal plot of all measured titers against homologous HA, homologous NA, and heterologous NA clearly shows adjuvanted PNP hydrogels increased the magnitude and breadth of antibody responses compared to commercial Fluzone (Figure 7d). A phylogenetic tree of the influenza surface proteins from the various strains examined in this study shows the genetic diversity of antigens toward which we have developed an immune response using the PNP-1-5/3M-052/Fluzone platform (Figure 7e).

**Figure 7.**
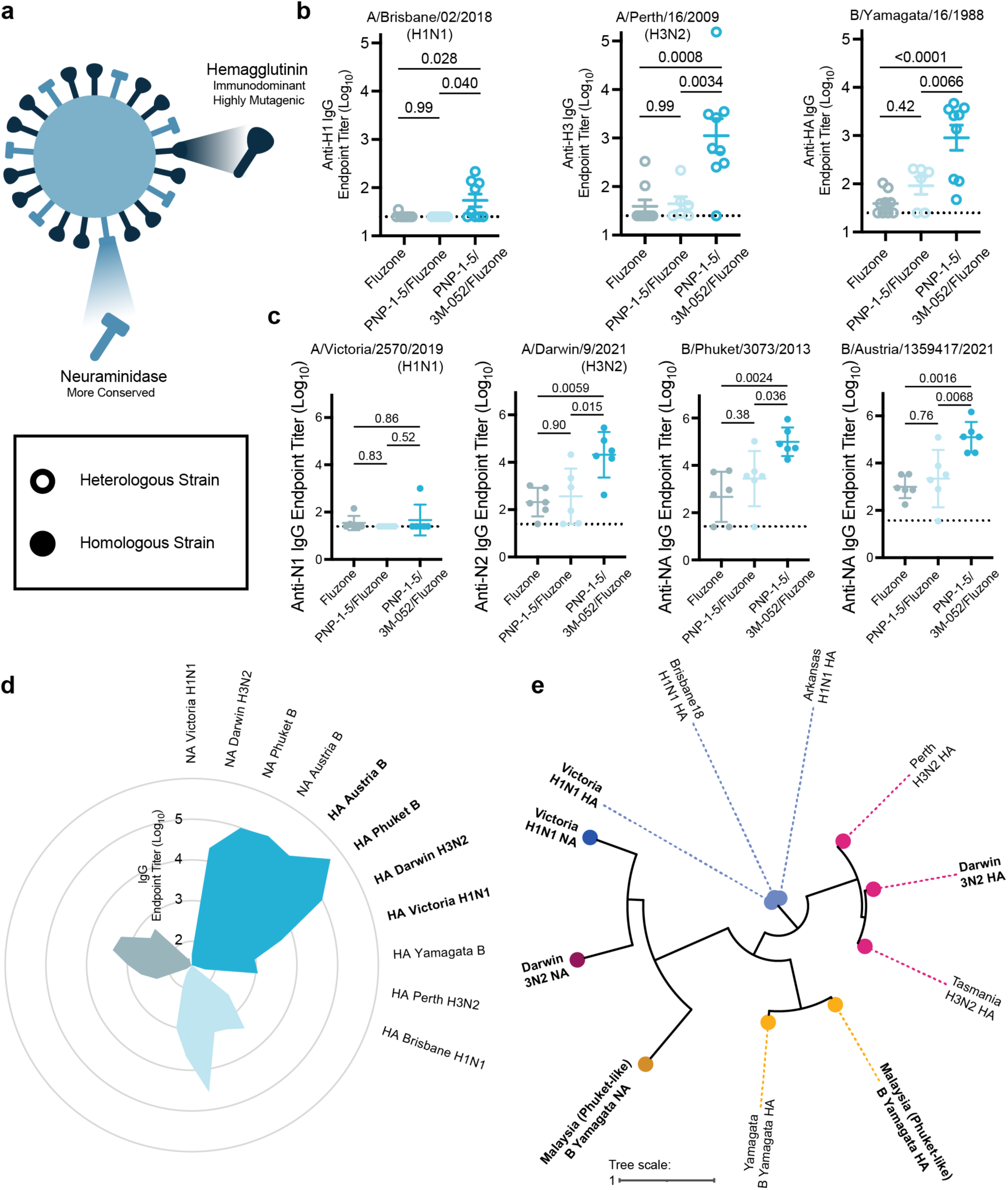
3M-052-adjuvanted PNP hydrogel vaccines provide the greatest breadth and response to viral activity. **a.** Diagram of two surface proteins, hemagglutinin and neuraminidase, on an influenza virus. **b.** Week 6 IgG titers against HA from heterologous viruses bearing the same HA subtype – H1, H3 – or type – B – as the HAs included in the administered vaccine. (n = 9) **c.** Week 8 IgG titers against NA of the vaccinated strains. (n = 6) **d.** Petal plot summarizing average IgG titers against all tested HAs and NAs (clockwise from top right: dark blue PNP-1-5/3M-052/Fluzone, light blue PNP-1-5/Fluzone, gray unadjuvanted Fluzone). **e.** Phylogenetic tree of A/Victoria/2570/2019 (H1N1), A/Darwin/9/2021 (H3N2), B/Malaysia/U2555/2013 (a B/Phuket/3073/2013-like virus), A/Brisbane/02/2018 (H1N1), A/Arkansas/08/2020 (H1N1), A/Perth/16/2009 (H3N2), A/Tasmania/503/2020 (H3N2), B/Yamagata/16/1988 (B Yamagata), influenza antigens based on HA or NA DNA sequence where the radial branch lengths represent modifications per site. Created using the ‘Viral Genome Tree’ tool on the Bacterial and Viral Bioinformatics Resource Center (BV-BRC) website https://www.bv-brc.org/ and formatted on https://itol.embl.de/. All error bars are mean ± SEM, P values determined by one-way ANOVA with Tukey’s multiple comparisons test. Statistical significance was considered as P < 0.05.

### 2.7 Enhanced Immunity to a Clinical Quadrivalent Vaccine

We then examined further metrics of vaccine efficacy to better understand the increase in breadth afforded by adjuvanted extended delivery vaccines. Hemagglutination inhibition (HAI) assays were performed on sera pooled across timepoints for each mouse using two homologous and two heterologous influenza strains (Figure 8a). The HA surface protein on influenza viruses causes agglutination of red blood cells and in turn prevents their settling out of solution. An HAI assay determines the concentrations of sera at which there is enough anti-HA antibody present to neutralize the HA of an influenza virus and prevent red blood cell agglutination, thereby allowing cells to settle out of solution. In all tested strains, PNP-1-5/3M-052/Fluzone generated the most virus inhibiting antibodies. Against the two homologous strains, PNP-1-5/Fluzone improves viral inhibition over commercial Fluzone, and PNP-1-5/3M-052/Fluzone improves even further with a significant increase in titer compared to Fluzone. This trend is echoed in evaluation of heterologous H1 virus, indicating breadth in HAI in addition to the general antibody magnitude breadth increase shown previously. A timeline of the vaccination for additional immune assays is shown in Figure 8b. At three weeks post-vaccination, an HA-specific IFNγ T cell ELISpot was performed on splenocytes from a cohort of mice (Figure 8c). Splenocytes were stimulated with an HA peptide pool from each homologous strain and IFNγ-producing cells were counted via spot detection. The resulting data showed comparable HA-specific IFNγ-producing T cell responses between commercial Fluzone and PNP-1-5/3M-052/Fluzone groups, indicating that prolonged antigen exposure with our PNP-hydrogel based vaccines does not impair T cell responses. Further, to investigate a possible cause of the increased antibody diversity, competitive ELISAs were performed 8 weeks after vaccination on polyclonal IgG purified from sera. Plates coated with A/Darwin/9/2021 H3N2 HA received an additional incubation with a constant concentration of one of two anti-influenza monoclonal antibodies (mAbs): (i) HA-stem-specific broadly neutralizing antibody (bnAb) MEDI8852 (MEDI), or (ii) HA-head-specific antibody FluA-20^[40]^ (Figure 8d). Competitive ELISA of these mAbs with a dilution series of mouse polyclonal IgG showed an order of magnitude difference between Fluzone and PNP-1-5/3M-052/Fluzone binding curves (Figure 8e,f). Remarkably, PNP-1-5/3M-052/Fluzone exhibited near identical binding in competition with MEDI or FluA-20 while commercial Fluzone showed significantly greater binding against MEDI (Figure 8g). In these assays, greater binding in competition with MEDI than FluA-20 means that the polyclonal antibody populations skew towards binding with the HA head region and are therefore less affected by the competitive stem-binding bnAb. This finding suggests that antibodies generated by sustained-release PNP hydrogel formulations elicit responses that are both higher affinity and more evenly split between head and stem binding, whereas IgG from commercial Fluzone vaccination is more subject to competition (i.e., lower affinity) and favors head binding.

**Figure 8.**
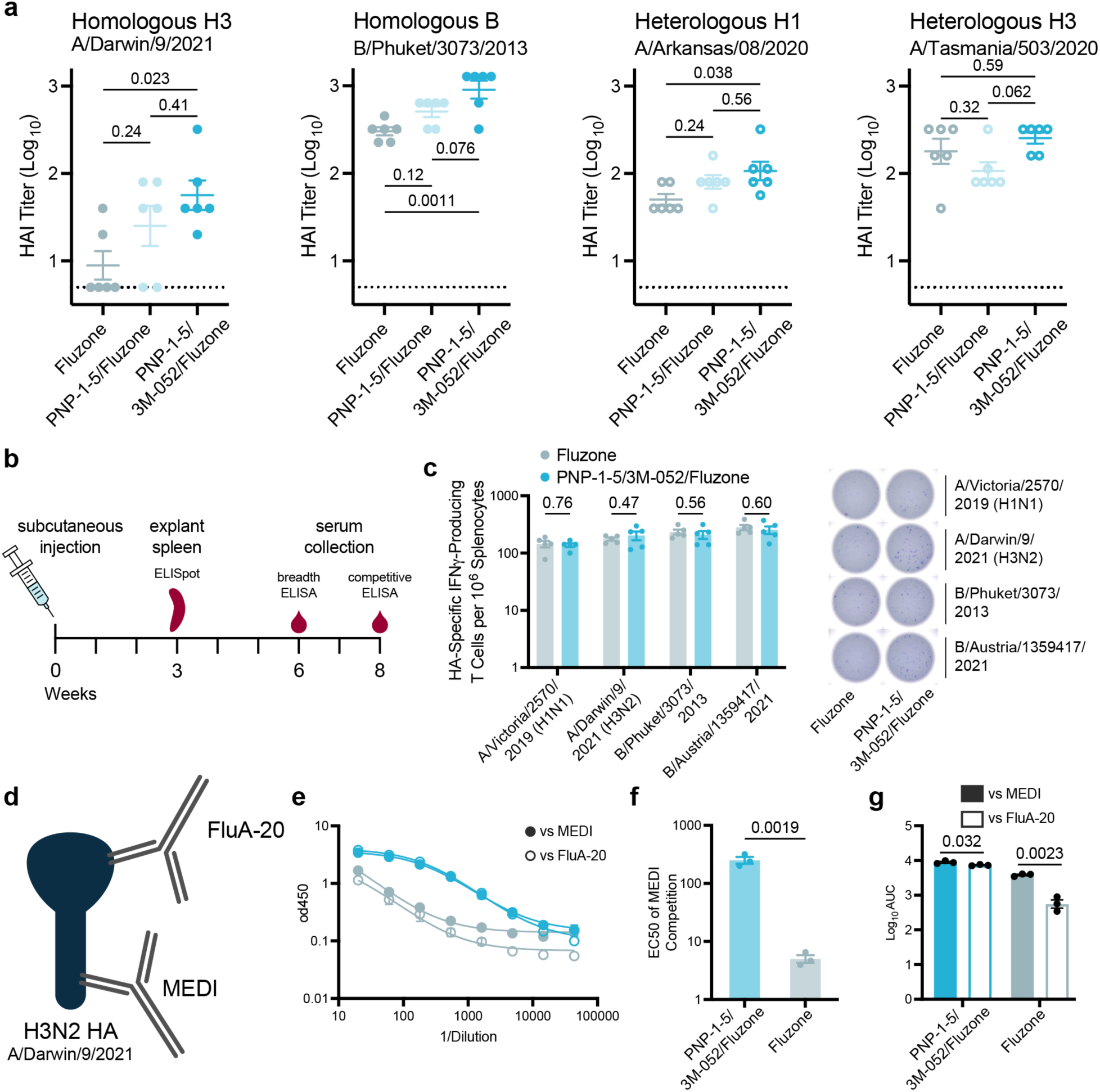
Mechanistic investigation into breadth of humoral response provided by adjuvanted PNP-hydrogel vaccines. **a.** Hemagglutination inhibition assay performed against homologous (H3 - A/Darwin/9/2021, B/Yamagata lineage - B/Phuket/3073/2013) or heterologous (H1 - A/Arkansas/08/2020, H3 - A/Tasmania/503/2020) influenza viruses on timepoint-pooled sera. (n = 6) **b.** Schedule of vaccination with Fluzone formulations as described in Figure 5 for additional immune assays. **c.** Homologous HA-specific IFNγ-producing T cells measured via splenocyte ELISpot at 3 weeks post vaccination. ELISpot well plate images show samples from one representative mouse in the Fluzone and PNP-1-5/3M-052/Fluzone groups in wells coated with each of the peptide antigens. (n = 6) **d.** Schematic of influenza mAbs FluA-20 and MEDI 8852 binding to head and stem regions, respectively, on HA. **e.** Competitive EILSA dilution curves of week 8 purified IgG against two mAbs, MEDI8852 (stem binder) and FluA-20 (head binder), binding to A/Darwin/9/2021 HA. (n = 3, replicates of sera pooled from 9 mice) **f.** EC50 of MEDI dilution curves in (e). **g.** Area under the curve (AUC) of dilution curves in (e). Dotted lines indicate lower limit of detection. All error bars are mean ± SEM, P values determined by one-way ANOVA with Tukey’s multiple comparisons test (a) or a two-tailed Student’s *t*-test (c, f, g). Statistical significance was considered as P < 0.05.

## 3. Discussion

Improving the influenza vaccine’s efficacy against both the strains included in the vaccine and against future viral variants is essential to preventing the roughly 500,000 deaths occurring annually and to safeguard against future pandemic outbreaks. Recent innovations in influenza vaccine technology such as mosaic virus-like particles or mRNA platforms are highly promising but require a complete reworking of the influenza vaccine production supply chain^[64]^. In this work we have sought to fabricate an adjuvanted hydrogel platform that can be easily used in conjunction with commercial multivalent subunit or inactivated virus vaccines to enhance the potency, durability, and breadth of humoral immune responses. Previous publications have shown that our PNP hydrogels are immunologically inert^[44]^, and that hydrogel depots administered subcutaneously only recruit immune cells when they comprise inflammatory cargo such as adjuvants^[33, 55]^. In this study we therefore sought to further enhance the immunogenicity of our extended-release platform with new molecular adjuvants. Our immunogenicity data indicate that sustained vaccine exposure with our PNP hydrogel technology ensures full seroconversion against all included viral strains, a development from our previous work which had only shown improvements in humoral immune responses with single antigen vaccines^[52–54, 56–58]^. The superior humoral immunity also extends to heterologous strains of influenza, where mice receiving hydrogel-based vaccines exhibited complete responses and comparable titers to those elicited by clinically relevant adjuvant technologies against the homologous strains. In contrast, many animals immunized with clinically relevant controls were unresponsive to these heterologous strains. We show that the observed enhancements in potency and breadth can be attributed to the production of a higher affinity and diverse array of effective antibodies, corroborating previous literature indicating that prolonged exposure of vaccines increases the magnitude and duration of germinal center reactions, increasing somatic hypermutation and commensurate affinity maturation^[33]^.

Our initial study focused on comparing HA-based subunit vaccines delivered either in liquid bolus vaccines with various clinically relevant adjuvants, including MF59-like and AS04-like systems, or in a PNP hydrogel system comprising either MPLA or CpG. Compared to Addavax (MF59-like), alum/MPLA (AS04-like), and alum/CpG adjuvanted vaccines, sustained vaccine exposure with adjuvanted PNP hydrogels resulted in a drastic improvement in IgG antibody production against all vaccinated strains, whereby all mice showed consistent high titers across the timepoints tested. In contrast, standard liquid bolus vaccine groups typically exhibited at least 20% non-responders to the homologous influenza strains incorporated in the vaccine. Effective seroconversion against administered strains varies widely in a clinical setting, with seroconversion rates among vaccinated adults often varying from 30-70% one month after vaccination^[65–67]^. The ability of PNP hydrogel based multivalent vaccines to induce consistently complete seroconversion is a marked improvement over current commercial vaccine technologies. Additionally, the dramatic improvement observed in titers elicited against heterologous HAs from various influenza subtypes and lineages moves towards addressing the fact that current influenza vaccines are largely ineffective against future mutations, even those which arise over the course of a typical flu season^[68]^. Hydrogel-based vaccines elicited improved titers against HAs from H1N1, H3N2, and B strains from both past and future years and different regions than the homologous strains, indicating improved protection against genetic variance as the viruses continually mutate. Furthermore, adjuvanted PNP hydrogels also elicited robust responses H5, H7, and H9 strains, indicating the potential of these vaccines to provide protection against zoonotic spillover of high-threat pandemic strains to which the human population has little to no protection^[8–10]^. These results with a subunit HA vaccine formulation demonstrated the ability of the PNP hydrogel depot platform to provide controlled HA delivery and potent immune stimulation to improve the consistency of seroconversion, potency, and breadth of a multivalent influenza vaccine.

We then sought to leverage adjuvanted PNP hydrogels as a simple-to-implement admix platform for improving immune responses to commercial Fluzone Quadrivalent. We again saw significant improvement in the consistency and potency of immune responses against both homologous and heterologous antigen responses. Additionally, PNP hydrogel-based vaccines elicited greatly enhanced hemagglutination inhibition titers to both homologous and heterologous viruses. While the Fluzone vaccine is made from split viruses and thereby comprises by HA and NA proteins, clinical studies typically only focus on anti-HA responses, even though the NA surface proteins are much more conserved as the virus mutates. Remarkably, we also found a significant enhancement in anti-NA titers for PNP hydrogel-based vaccines. To evaluate the molecular basis for the increased antibody breadth, we conducted competitive binding assays with broadly neutralizing MEDI (stem-targeted) and FluA-20 (head-targeted) mAbs. These studies revealed that while conventional Fluzone vaccines elicit IgG responses that are primarily head-binding and rather low affinity, adjuvanted PNP hydrogel vaccines induce a more balanced response between head and stem binding antibodies and much higher affinity, overcoming the typical immunodominance of the head-binding region of the HA through prolonged exposure. It has previously been postulated that directing immune response towards the more conserved stem region of the HA could gain better year-to-year immunity^[69–70]^. The combination of more HA stem-targeted responses and NA responses indicates that the PNP hydrogel-based vaccines my provide much broader protection.

Overall, a universal influenza vaccine must be scalable, potent against current strains, and robust to mutations of the virus^[71]^. The extensive antibody range induced by the adjuvanted PNP hydrogel platform against varied strains and subtypes, poorly immunogenic NA antigens, as well as historically difficult HA antigen regions is evidence that sustained delivery can increase the diversity and potency of humoral responses to influenza vaccination^[20]^. A breadth of anti-HA antibodies is known to be protective against infection by the virus^[18, 72]^ and furthermore increases the chance of gaining immunity that will be maintained against future mutations. The ease of use of our adjuvanted PNP hydrogel system, shown in this work to readily admix with commercial Fluzone, provides a promising way to “supercharge” current influenza vaccination infrastructure to better protect our global populations against both annual influenza and pandemic risks.

## 4. Conclusion

Through these studies, we show the development of an easy-to-implement adjuvant system for enhancing existing multivalent influenza vaccines comprising a self-assembled, depot-forming, adjuvanted PNP hydrogel system capable of eliciting broad, potent, and durable humoral responses. In comparison to standard liquid bolus vaccines, PNP-hydrogel based vaccines induced higher antibody titers, more complete seroconversion, higher hemagglutination inhibition, and broader responses to heterologous strains of influenza, including several pandemic threats. The breadth of protection can be attributed to a greater diversity of antibody binding sites, which allow binding to more conserved regions of HA proteins. This platform has the potential to significantly improve our preparedness for a potential influenza pandemic.

## 5. Materials and Methods

### Materials

HPMC (US Pharmacopeia Grade), N,N-Diisopropylethylamine (Hunig’s base), hexanes, diethylether, 1-dodecyl isocyanate (99%), N-methyl-2-pyrrolidone (NMP), dichloromethane (DCM), 3,6-dimethyl-1,4-dioxane-2,5-dione (lactide), 1,8-diazabicyclo(5.4.0)undec-7-ene (DBU, 98%), acetonitrile (ACN), dimethyl sulfoxide (DMSO), Poly(ethylene glycol)–methyl ether (PEG, 5 kDa), bovine serum albumin (BSA) were purchased from Sigma Aldrich and used as received. CpG1826 (Vac-1826), MPLAs (Vaccigrade), Addavax, and alum (Alhydrogel 2%) were purchased from InvivoGen. 3M-052 was purchased from 3M and the Access to Advanced Health Institute (AAHI). AF-647 Succinimidyl Ester was purchased from Thermo Fisher Scientific. Influenza hemagglutinin and neuraminidase proteins were purchased from Sino Biological: H1N1 (A/California/07/2009) HA, H3N2 (A/Perth/16/2009) HA, B (B/Brisbane/60/2008) HA, H3N2 (A/Indiana/07/2012) HA, H3N2 (A/Hong Kong/CUHK31987/2011) HA, B (B/Yamagata/16/1988) HA, H1N2 (A/Minnesota/14/2012) HA, H1N1 (A/Brisbane/02/2018) HA, H1N1 (A/Brisbane/59/2007) HA, H5N1 (A/Bangladesh/207095/2008) HA, H7N9 (A/Shanghai/1/2013) HA, H9N2 (A/Hong Kong/1073/99) HA, H1N1 (A/Victoria/2570/2019) HA and NA, B (B/Phuket /3073/2013) HA and NA, B (B/Austria/1359417/2021) HA and NA, H3N2 (A/Darwin/9/2021) HA and NA. PepMix of HA/H1N1/Victoria/2019, HA/H3N2/Darwin/2021, HA/H0N0/Phuket/2013, and HA/H0N0/Austria/2021 were purchased from JPT. Fluzone Quadrivalent and Fluzone High-Dose Quadrivalent were purchased from the Stanford pharmacy. FluA-20 and MEDI8852 mAbs were provided by the Kim lab at Stanford^[40]^. Goat anti-mouse IgG Fc secondary antibody (A16084) horseradish peroxidase (HRP) was purchased from Invitrogen and 3,3″,5,5″-Tetramethylbenzidine (TMB) ELISA substrate, high sensitivity, was purchased from Abcam.

### Preparation of HPMC-C_12_

HPMC-C_12_ was prepared according to previously reported procedures, and the protocols will be briefly described here. HPMC (1.0g) was dissolved in NMP (40mL) by stirring at room temperature overnight. The solution was then heated to 50°C for 45 minutes while 1-dodecyl isocyanate (125μL) was dissolved in NMP (5.0 mL). This solution was added dropwise to the reaction mixture, followed by ∼10 drops of Hunig’s base (catalyst). The reaction mixture was stirred at room temperature for 18 h. This solution was then precipitated from acetone, decanted, re-dissolved in water (∼2 wt%) and placed in a dialysis tube for dialysis against water for 3-4 days. The polymer was lyophilized and reconstituted to a 60 mg/mL solution with sterile PBS.

### Preparation of PEG-PLA

PEG-PLA was prepared as previously reported, and the protocols will be briefly described here. PEG (5kDa; 2.5g) was melted at to 90°C and dried on vacuum for 30 minutes. Recrystallized lactide (10g) was measured into a 250 mL round bottom flask and flushed with nitrogen. Dry, distilled DCM (50mL) was added to the flask to dissolve the lactide with light heating. DCM (5mL) was added to the melted peg to dissolve, followed by DBU (75μL, 0.1mmol; 1.4 mol% relative to lactide). The PEG solution was added rapidly to the lactide solution and was allowed to stir for 8 min. The reaction mixture was quenched by opening to air and precipitated from a 1:1 solution of hexane and diethylether. The synthesized PEG-PLA was collected and dried under vacuum. Gel permeation chromatography (GPC) was used to verify that the molecular weight and dispersity of polymers meet our quality control (QC) parameters.

### Preparation of PEG-PLA NPs

NPs were prepared as previously reported. A 1mL solution of PEG-PLA in 75:25 Acetonitrile:DMSO mixture (50mg/mL) was added dropwise to 10mL of water at room temperature under a high stir rate (600rpm). NPs were purified by ultracentrifugation over a filter (molecular weight cut-off of 10kDa; Millipore Amicon Ultra-15) followed by resuspension in PBS to a final concentration of 200 mg/mL. NPs were characterized by dynamic light scattering (DLS) to find the NP diameter (PEG-PLA NPs, 33 ± 3 nm).

### General PNP Hydrogel Preparation

The PNP-2-10 hydrogel formulation contained 2 wt% HPMC-C_12_ and 10 wt% PEG-PLA NPs in PBS. These gels were made by mixing a 2:3:1 weight ratio of 6 wt% HPMC-C_12_ polymer solution, 20 wt% NP solution, and PBS. The PNP-1-5 hydrogel formulation contained 1 wt% HPMC-C_12_ and 5 wt% PEG-PLA NPs in PBS. These gels were made by mixing a 2:3:7 weight ratio of 6 wt% HPMC-C_12_ polymer solution, 20 wt% NP solution, and PBS.

### Rheological Characterization

Rheological characterization was performed on a TA Instruments Discovery HR-2 torque-controlled rheometer with Peltier stage using a 200-m serrated plate geometry. Oscillatory amplitude sweeps were performed from 1% to 10,000% strain. Dynamic oscillatory frequency sweep measurements were performed with a constant strain of 1% from 0.1 rad/s to 100 rad/s. Steady shear experiments were performed from 100 to 0.1 s^-1^.

### Injection Force Measurements

Force of injection was quantified by measuring the required force to inject a PNP gel through a known needle gauge at a flow rate of 2 mL per minute using a 1 mL syringe of known barrel dimensions. A force sensor was built that encompassed a load cell (FUTEK LLB300 50 lb Subminiture Load Button (Model #: LLB300, Item #: FSH03954, Serial #: 705242) attached to a syringe pump (KD Scientific Syringe Pump (Model #: LEGATO 100, Catalog #: 788100, Serial #: D103954)). An Omega Engineering Platinum Series Meter (Model #: DP8PT, Serial #: 18110196) was used to translate load cell resistance measurements to force values in kg. The load cell was calibrated prior to measuring injection force. A lab view program records the forces measured throughout the duration of an injection experiment and displays a graph of injection force over time. Injection force experiments were performed as follows. A 1mL Thermo Fisher luer lock syringe equipped with a 27 gauge ultra-thin-wall, ½ inch (TSK SteriJect PRC-270131-100) needle was loaded into the syringe pump. The syringe pump height was adjusted so that the load button of the force sensor was in contact with the end of the syringe plunger. The initial force was at or very close to 0 kg. The appropriate syringe barrel dimensions as well as desired flow rate of 2 mL per minute and injection volume of 200 μL were then selected. The syringe pump moved at the programmed rate injecting PNP hydrogel through the attached needle. The force sensor coupled with the Omega unit measured the force required to inject the PNP hydrogel at the desired flow rate. A lab view program recorded the forces measured throughout the duration of an injection experiment and displayed a graph of injection force over time. Force of injection was quantified by subtracting the average initial force (background) from the average plateau injection force. Injection force in kg was converted to injection force in Newtons by multiplying by 9.81.

### Cargo Release In Vitro

Glass capillary tubes capped with epoxy at one end were incubated for 1 hour with a 1% BSA in 1x PBS solution for 1 hour and then dried overnight. PNP hydrogel containing the cargo of interest (100μL) was injected into the bottom each tube and then covered with 400μL 1x PBS. Tubes were covered and incubated at 37°C between measurement timepoints. The 400μL of PBS was removed via syringe without disturbing the gel at each timepoint and measured for fluorescence to determine mass released using a Synergy H1 Microplate Reader (BioTek Instruments). Fresh 400μL PBS was immediately replaced on top of the gel.

### Alexa Fluor 647 Conjugated Hemagglutinin Protein

A premixed solution of AF-647 Succinimidyl Ester (30 μg, 0.028 μmol, 9 equiv, 5mg/mL stock solution in DMSO) in PBS 1X was added to a solution of hemagglutinin protein (H3N2 (A/Perth/16/2009) HA, 200 μg, 0.18 μmol, 1 equiv) in PBS 1X. A volume ratio of dye (1/10) to protein (9/10) was respected. The reaction was conducted in the dark for 4 h at RT with mild shaking. The solution was quenched by diluting 2-fold with PBS 1X and purified in centrifugal filters (Amicon Ultra, 10 kDa MWCO 0.5 mL) at 14g for 10 min. The purification step was repeated until all excess dye was removed. The solution was then resuspended in PBS 1X and stored at −20 °C.

### In Vivo Pharmacokinetic Study of HA Protein in Bolus and Hydrogel Formulations

SKH1E mice were immunized subcutaneously in the right flank with 100 μL of liquid bolus or hydrogel vaccines containing 5 μg of AF647-HA protein and 10 μg of MPLAs. Hydrogels were formulated to a 2-10 formulation as described in previous sections. Bolus liquid administration contained 75 μg alhydrogel. Mice were imaged over 25 days using an *in vivo* imaging system (IVIS Lago). AF647-HA proteins were imaged using an auto exposure time, an excitation wavelength of 640 nm, and an emission wavelength of 710 nm (binning: medium, F/stop: 2). Average radiant efficiency was quantified and used to calculate area under the curve. Data were fitted with a one-phase decay model in Prism to calculate time to 80% release, t_80%_. Data sets for which Prism could not generate a 95% confidence interval for the fit were instead measured using the “cut off” method and assigned the value of the last day at which the signal was above the 80% release value. Data analysis was performed by using GraphPad Prism.

### Vaccine Formulations Comprising Trivalent Subunit

Injections volumes of 100 μL were prepared for each dose. Influenza vaccines contained 5μg of each hemagglutinin (HA) from each of 3 influenza strains: Influenza A H1N1 (A/California/04/2009), Influenza A H3N2 (A/Perth/16/2009), and Influenza B (B/Brisbane/60/2008) (Sino Biological). Experimental groups were adjuvanted with the following: (liquid) 1625μg Addavax, (liquid) 20μg CpG + 75μg alum, (liquid) 20μg MPLAs + 75μg alum, (gel) 20μg CpG + PNP-2-10 hydrogel, (gel) 20μg MPLAs + PNP-2-10 hydrogel. PNP-2-10 contained a ratio of 2 wt% HPMC-C_12_ : 10 wt% NP. For the PNP hydrogels, the vaccine cargo was added at the appropriate concentration into the PBS component of the hydrogel before adding the polymer and NP solutions, as described above.

### Vaccine Formulations Comprising Commercial Fluzone

Injections volumes of 200μL were prepared for each dose. The influenza vaccine contained diluted amounts of 2022-2023 Fluzone Quadrivalent (liquid formulation) or Fluzone High-Dose Quadrivalent (gel formulations) dosed to provide 2.5μg of each hemagglutinin (HA) from each of the 4 included influenza strains: Influenza A H1N1 (A/Victoria/2570/2019), Influenza A H3N2 (A/Darwin/9/2021), Influenza B (B/Phuket /3073/2013), Influenza B (B/Austria/1359417/2021). Experimental groups were adjuvanted with the following: (liquid) nothing, (gel) PNP-1-5, (gel) 1μg 3M-052 + PNP-1-5, (gel) 10μg MPLAs + PNP-1-5. PNP-1-5 contained a ratio of 1 wt% HPMC-C_12_ : 5 wt% NP. For the PNP hydrogels, the vaccine cargo was added at the appropriate concentration into the PBS component of the hydrogel before adding the polymer and NP solutions, as described above.

### Animal Protocols

NIH guidelines for the care and use of laboratory animals (NIH Publication #85-23 Rev. 1985) have been observed. All animal studies were performed with the approval of Stanford Administrative Panel on Laboratory Animal Care.

### Vaccination

C57BL/6 mice were purchased from Charles River and housed at Stanford University. Female mice between 6 and 10 weeks of age at the start of the experiment were used. The mice were shaved several days before vaccine administration and received a subcutaneous injection (100μL or 200μL administration volume) of gel or liquid vaccine on their backs under brief isoflurane anesthesia. Blood was collected from the tail vein at predetermined timepoints for survival studies.

### Mouse Serum ELISAs to Determine Antibody Concentration

Serum IgG antibody titers for the influenza vaccine were measure using an ELISA. Maxisorp plates (Thermofisher) were coated with HA or NA (Sino Biological) at 4μg/mL in PBS overnight at 4°C and then blocked with PBS containing 1% non-fat dry milk for 1 h at 25°C. Serum was diluted into a 1% BSA in PBS solution in a v-bottom plate at 1:100 and then four-fold serial dilutions were performed up to 1:1,638,400 dilution. Titrations were added to plates and after 2h at 25°C, goat– anti-mouse IgG Fc-HRP (1:10,000, Invitrogen, A16084) was added for 1h at 25°C. Plates were developed with TMB substrate (TMB ELISA Substrate (High Sensitivity), Abcam). The reaction was stopped with 1 M HCl. The plates were analyzed using a Synergy H1 Microplate Reader (BioTek Instruments) at 450nm. Dilution curves were fitted to an inhibitor vs. response three parameter curve and endpoint titers were defined as the reciprocal of the serum dilution that gave an optical density of 0.1 (twice the limit of detection) on the fitted curve.

### Competitive ELISA

IgG purified from the D56 sera of Fluzone and PNP-1-5/3M-052/Fluzone groups was used to perform a competitive ELISA. IgG from the same experimental group was pooled and experiments were done in triplicate. ELISA was performed as reported above with the following modifications: After coating plate with A/Darwin/9/2021 HA and blocking, an additional step was added: incubation with human FluA-20 or MEDI8852 mAbs at 50μg/mL for 30 minutes. IgG dilutions were then applied: dilutions started at 1:20 and then three-fold serial dilutions were performed up to 1:43,740 dilution. Goat–anti-mouse IgG Fc-HRP was added as before to detect the mouse IgG without interference from the human mAbs and development, stopping, and readout were performed as written above.

### ELISpot

The frequency of antigen-specific IFN-γ producing splenocytes was determined by measuring antigen-specific IFN-γ producing CD8+ T cells using a Mouse IFN-γ Single-Color ELISPOT kit (CTL ImmunoSpot). Spleen cells were harvested 3 weeks post-vaccination following immunization with Fluzone groups. 400,000 splenocytes per sample were pipetted into the well and were stimulated with 1μg/mL a mixture of peptides from each of the 4 distinct antigens (JPT) for 24h at 37°C. Spots were then developed following manufacturer’s instruction (Immunospot S6 Ultra M2).

### Statistical Analysis

Comparisons between two groups were conducted by a two-tailed Student’s t-test. One-way ANOVA test with a Tukey’s multiple comparisons test was used for comparison across multiple groups. Statistical analysis was run using GraphPad Prism 10 (GraphPad Software). Statistical significance was considered as P < 0.05. Data in plots were represented as mean ± SEM.

## 6. Data Availability Statement

The data that support the findings of this study are available from the corresponding author upon reasonable request. A preprint of this work is available through bioRxiv.org^[73]^.

## Acknowledgements

This work was supported in part by the Bill & Melinda Gates Foundation (INV-027411). O.M.S., J.Y., and C.K.J. are thankful for a National Science Foundation Graduate Research Fellowship. O.M.S. is also thankful for Hancock Fellowship of the Stanford Graduate Fellowship in Science and Engineering. B.S.O. is grateful for an Eastman Kodak Fellowship. We thank Dr. Duo Xu in the laboratory of Dr. Peter Kim for providing mAbs. We extend our gratitude to Dr. Thomas Oguin at the Duke Virology RBL for performing HAI assays. We also would like to thank the staff of the BioE/ChemE Animal Facility who cared for the mice.

