## Supplementary Materials for "Sustained Vaccine Exposure Elicits More Rapid, Consistent, and Broad Humoral Immune Responses to Multivalent Influenza Vaccines"

**The PDF file includes:**

Figs. S1 to S5

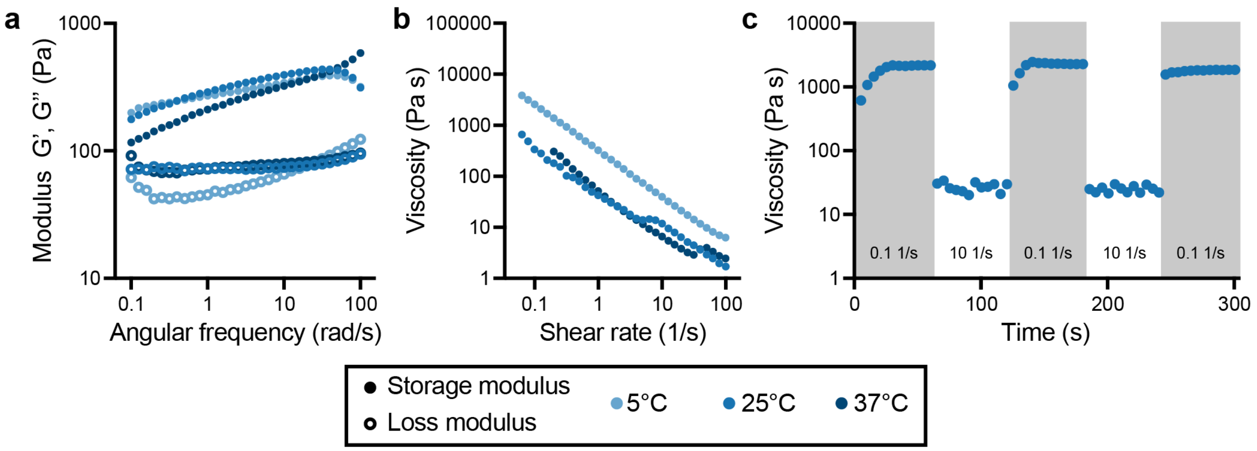


**Fig. S1. Rheological tests performed on a PNP-2-10 hydrogel.**
**a.** Data from frequency-dependent oscillatory shear experiments at three relevant temperatures indicates that the PNP hydrogel materials exhibit solid-like properties with the storage and loss modulus (G' and G", respectively) that are invariant to temperatures relevant for storage, administration, and in vivo efficacy, respectively (evaluated at 1% strain). **b.** Shear-dependent rheological data from a steady shear rate sweep at three temperatures shows robust shear-thinning behaviors that are maintained across all temperatures evaluated. **c.** Step-shear measurements in which the hydrogels were subjected to cycles of low and high shear rates ($0.1\ s^{-1}$, $10\ s^{-1}$) in 60 s steps at 25°C demonstrate the hydrogel's ability to shear-thin and rapidly self-heal under flow conditions mimicking the process of injection.

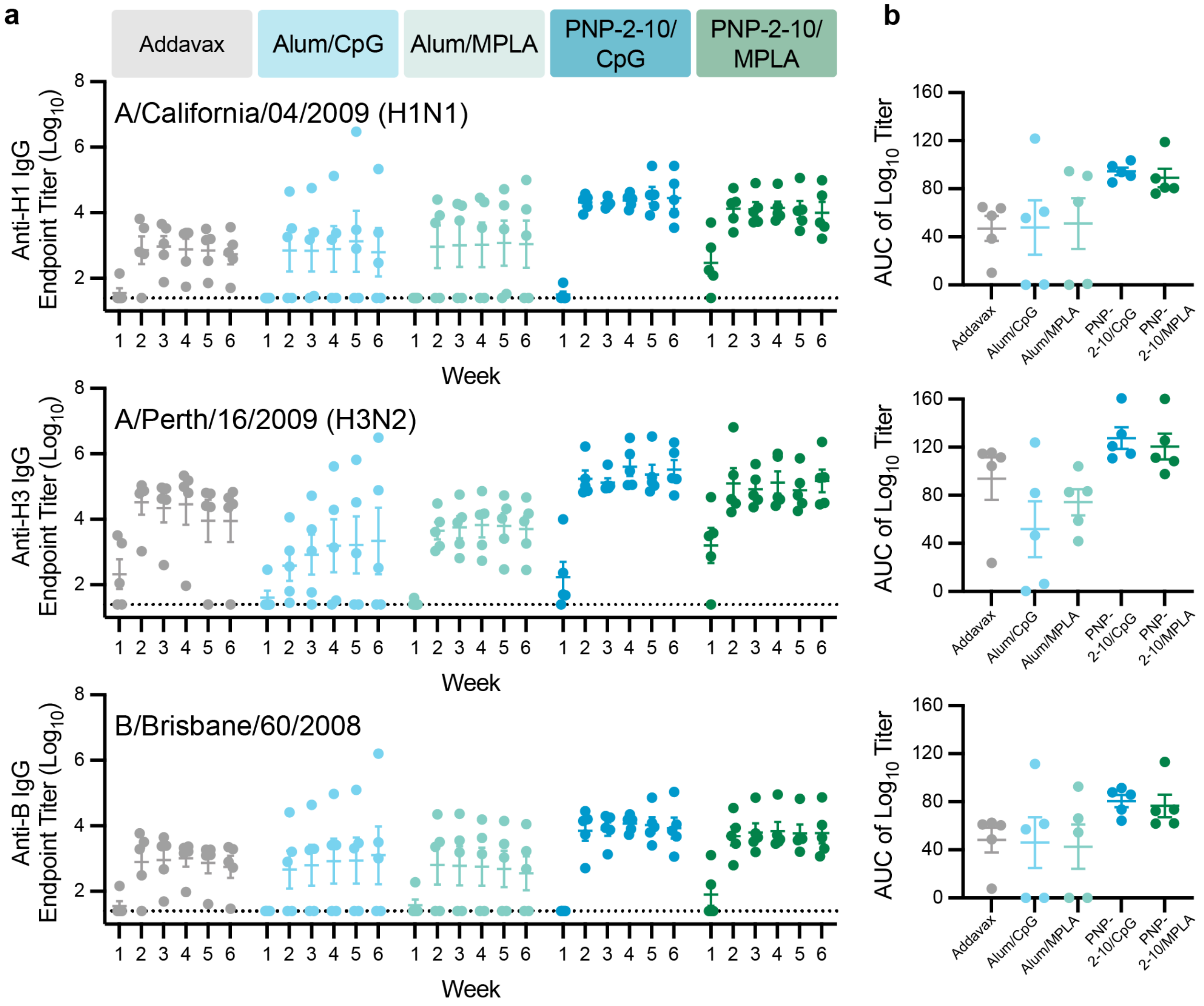


**Fig. S2. Trivalent subunit influenza vaccine adjuvanted by PNP hydrogels.**
Sustained delivery of multivalent flu vaccines induces robust, consistent humoral responses against all homologous strains. Mice (C57BL/6) were immunized with a single 100 µL dose of trivalent influenza vaccine containing 5 µg each of three hemagglutinin proteins (HA) - Influenza A H1N1 (A/California/07/2009), Influenza A H3N2 (A/Perth/16/2009), Influenza B (B/Brisbane/60/2008) in one of five different vaccination strategies – Addavax, Alum/ CpG, Alum/MPLA, PNP-2-10/CpG, PNP-2-10/ MPLA. (n = 5). **a.** Anti-HA IgG titers from vaccinated mice over 6 weeks. Dotted lines indicate lower limit of detection. **b.** Area under the curve generated from the titers shown in part (a).

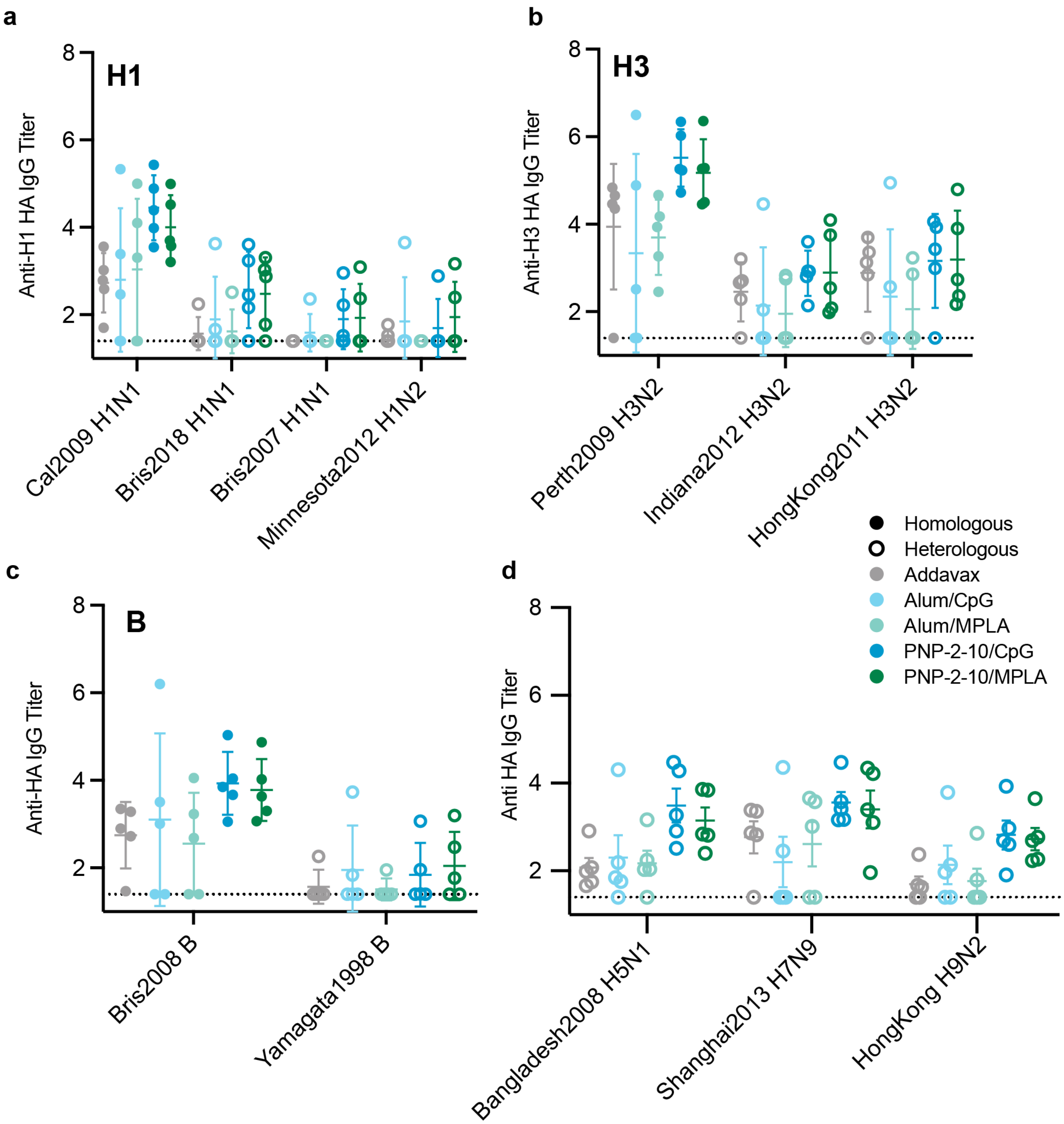


**Fig. S3. Breadth of HA titers from trivalent vaccination against heterologous strains.** Week 6 IgG titers against HA from heterologous viruses bearing the same HA subtype or type, including **a.** H1, **b.** H3, and **c.** B, as the HAs included in the administered vaccine. **d.** Week 6 anti-HA IgG titers against example H5, H7, and H9 viruses, which are viral strains of different subtypes than the HAs included in the administered vaccine. Dotted lines indicate lower limit of detection.

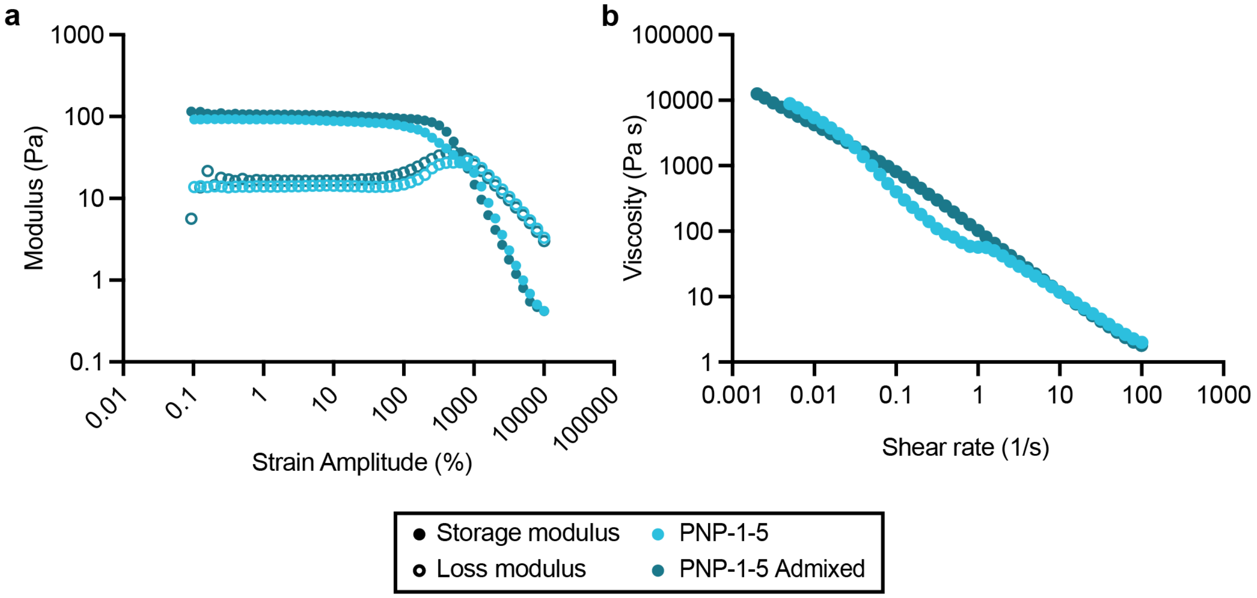


**Fig. S4. Rheology of PNP-1-5 standard and admixed formulations.**
**a.** Stress-controlled oscillatory amplitude sweep rheological data of PNP-1-5 hydrogels with different preparation procedures. **b.** Shear-dependent rheological data from a steady shear rate sweep shows decreasing viscosity with increasing shear rate for PNP-1-5 hydrogels prepared with different procedures.

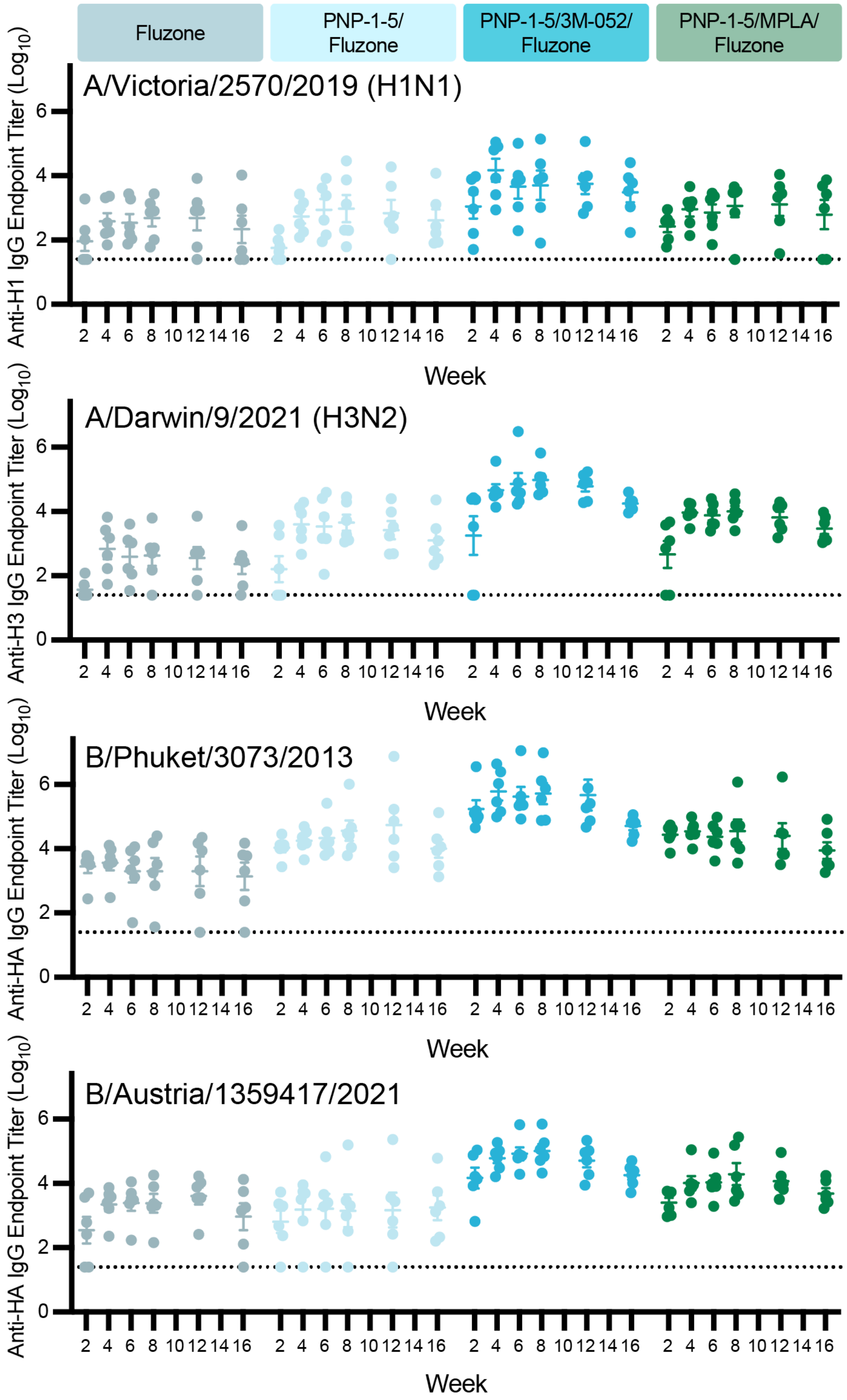


**Fig. S5. Homologous anti-HA titers from clinical Fluzone Quadrivalent vaccination.**
In this experiment, mice were immunized with vaccines comprising 2.5 μg HA protein from each strain contained within the 2022-2023 Fluzone Quadrivalent vaccine and an optional adjuvant (e.g., commercial control, PNP-1-5, PNP-1-5/3M-052, PNP-1-5/MPLA) (n = 6). Anti-HA IgG titers from vaccinated mice over 16 weeks. Dotted lines indicate lower limit of detection.
